# CIRFESS: An interactive resource for querying the set of theoretically detectable peptides for cell surface and extracellular enrichment proteomic studies

**DOI:** 10.1101/2020.01.22.916148

**Authors:** Matthew Waas, Jack Littrell, Rebekah L. Gundry

**Affiliations:** CardiOmics Program, Center for Heart and Vascular Research; Division of Cardiovascular Medicine; and Department of Cellular and Integrative Physiology, University of Nebraska Medical Center, Omaha, NE, 68198, USA

## Abstract

Cell surface transmembrane, extracellular, and secreted proteins are high value targets for immunophenotyping, drug development, and studies related to intercellular communication in health and disease. As the number of specific and validated affinity reagents that target this subproteome are limited, mass spectrometry (MS)-based approaches will continue to play a critical role in enabling discovery and quantitation of these molecules. Given the technical considerations that make MS-based cell surface proteome studies uniquely challenging, it can be difficult to select an appropriate experimental approach. To this end, we have integrated multiple prediction strategies and annotations into a single online resource, <u>C</u>ompiled <u>I</u>nteractive <u>R</u>esource <u>f</u>or <u>E</u>xtracellular and <u>S</u>urface <u>S</u>tudies (CIRFESS). CIRFESS enables rapid interrogation of the human proteome to reveal the cell surface proteome theoretically detectable by current approaches and highlights where current prediction strategies provide concordant and discordant information. We applied CIRFESS to identify the percentage of various subsets of the proteome which are expected to be captured by targeted enrichment strategies, including two established methods and one that is possible but not yet demonstrated. These results will inform the selection of available proteomic strategies and development of new strategies to enhance coverage of the cell surface and extracellular proteome. CIRFESS is available at www.cellsurfer.net/cirfess.

## Introduction

The emergence of proteomics as a major discipline within the life science has been in no small part due to the development and eager adoption of computational strategies to enable the rapid analysis of mass spectrometry (MS) data files and inferred biological results. Since 1994, when the Yates laboratory introduced SEQUEST^1^, the first computational tool for fully automated database searching, continued developments in database construction, algorithm design, and software development have propelled the evolution of MS-based proteomics^2–9^. All aspects of MS-based proteomics, including interpretation of raw spectra and database searching, visualization of results, and subsequent biological inferences benefit from advances in bioinformatics. Beyond the analysis of experimental data, data science tools that integrate machine-learning or ontological resources have become increasingly popular for prediction and classification of protein-level information, a subject of recent review^10^. Such approaches rely on experimental data to train or inform predictions and annotations, and in turn, the prediction strategies can benefit experimental design.

To scientists at the bench, perhaps the most exciting and impactful bioinformatic tools are those that can inform the next experiment. To this end, web-based formats have become increasingly popular resources as they often require less setup (*e.g.* installation), avoid operating system compatibility issues, and can be used in a familiar framework. Hundreds of web-based bioinformatics tools are now available for proteomics (*e.g.* www.expasy.org/proteomics). Current tools span a broad range of utility, including systems-level distribution of proteins based on experimental observations, visualization of experimental results, and prediction and cataloging of specific post-translational modifications, interactomes, and subcellular proteomes. Despite the increase in availability of web-based proteomics tools, there are currently relatively few resources designed to specifically assist in experimental design and analysis of the cell surface proteome.

The cell surface and extracellular space contain proteins which play key roles in a wide range of biological processes and can be utilized as valuable markers for immunophenotyping and drug targets. Despite their importance, the cell surface proteome remains relatively poorly characterized compared to the depth that most intracellular proteomes have been described. Given the relative low abundance, presence of hydrophobic transmembrane spanning regions, and dynamic nature of the cell surface proteome owing to continuous cycling of proteins due to internalization, secretion, and stimulus-triggered recruitment to the plasma membrane, specialized techniques are typically required to enhance the detection of cell surface proteins by MS. Such proteomic approaches include enrichment strategies which exploit the biophysical properties of membranes - such as density gradient flotation, differential centrifugation, or silica-bead capture^11–15^ - or affinity-based approaches that use proximity labels^16–18^, lectins^19^, metabolic^20–22^ or chemical labels^23–27^ to enrich cell surface proteins. Application of these approaches have supported efforts to catalog the cell surface and secretome and have led to large scale efforts in experimentation^28^ and collation^29^. As with most proteomic methods, currently available strategies to probe the cell surface and secreted proteome are biased towards proteins that contain specific features *(e.g.* presence of an *N-*glycosylation site or lysine within an extracellular region of the protein that will generate a detectable peptide after trypsin digestion). Also, the implementation of these approaches can be inconsistent among users, resulting in variability in the specificity (*i.e.* cell surface versus non-cell surface) of enrichment observed. Hence, the integration of bioinformatic predictions with experimental data provides orthogonal means to interpret and filter results. Relevant predictions for surface and extracellular proteins include the presence of signal peptides^30–36^ and transmembrane domains^30,31,37–39^. Other approaches have applied manual curation, ontological annotations, or machine learning approaches to predict the subset of proteins that are localized to the cell surface and extracellular regions^40–43^. However, not all cell surface proteins contain canonical signal peptides^44^. Also, GPI-anchored and extracellular matrix or secreted proteins do not contain transmembrane domains, and gene ontology annotations may be insufficiently specific *(e.g.* cell surface versus membrane). Thus, filtering a proteomic dataset by these constraints often does not provide the complete picture of the cell surface proteome for a specific cell type.

Based on these limitations, in theory, it remains necessary to rely on experimental data to precisely define the proteins localized to the cell surface in a specific cell type. This leads to the question: *Which experimental approach is the best to use?* No doubt, the answer will be context dependent. If a specific monoclonal antibody is available, flow cytometry can be an effective approach for determining the surface localization of a protein. If antibodies are not available, the MS-based proteomic method of choice will depend on whether a cell surface proteome-wide screen or detection of a particular protein or subclass of proteins is desired. It will depend on the type and availability of the source material (*e.g.* metabolic labeling approaches cannot be used routinely for the analysis of primary human cells). Given the numerous technical considerations that make cell surface proteome studies uniquely challenging, it can be difficult to decide which approach to use. Currently, there is no single bioinformatic tool that can assist the investigator in determining which MS-based method is likely to be the most suitable approach for surface proteome studies.

To address this, we constructed a resource that integrates multiple prediction strategies and annotations relevant for the analysis of cell surface and extracellular proteins by MS and applied it to interrogate the human proteome. The results from these resources were compiled into a single interface and are accessible via a web-application termed <u>C</u>ompiled <u>I</u>nteractive <u>R</u>esource <u>f</u>or <u>E</u>xtracellular and <u>S</u>urface <u>S</u>tudies (CIRFESS), accessible at www.cellsurfer.net/cirfess. By bringing together key resources used to interrogate the surface and extracellular space, CIRFESS helps to prevent duplication of efforts and continued pinging of separate prediction servers for the same set of proteins. We expect CIRFESS will be informative for a broad range of future applications and will inform the selection or development of proteomic strategies to enhance coverage of the cell surface and extracellular proteome.

## Methods

### Database and prediction server access

The human reference proteome was downloaded from UniProt (canonical only, 20416 entries, accessed Sept 13, 2019). The proteome was filtered and split to meet the requirements of the individual prediction servers (*e.g.* length of proteins, number of entries). Default scoring settings were applied, and the outputs were collected as specified: TMHMM– ‘one line per protein’, Phobius – ‘Short’, PrediSi, – ‘Text’, Signal P – ‘Short output’. For analysis involving the protein-level evidence, the “Protein existence” field for each accession number was retrieved from UniProt. To aid in interpretation, a summary of the different categories is provided in the Table 1 (adapted from https://www.uniprot.org/help/protein_existence).

**Table 1:**
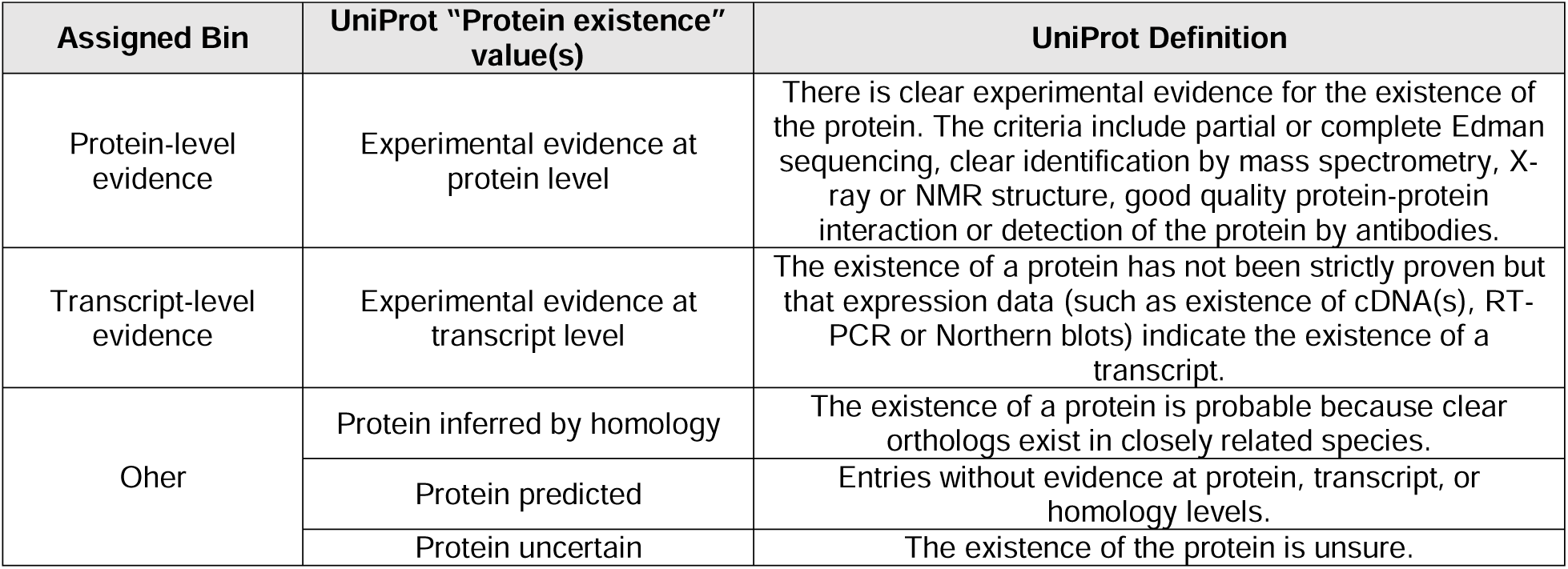
Summary of UniProt levels of “Protein existence” and the corresponding bins used in the analysis of levels of evidence for the different subsets of proteins.

| Assigned Bin | UniProt “Protein existence” value(s) | UniProt Definition |
| --- | --- | --- |
| Protein-level evidence | Experimental evidence at protein level | There is clear experimental evidence for the existence of the protein. The criteria include partial or complete Edman sequencing, clear identification by mass spectrometry, X-ray or NMR structure, good quality protein-protein interaction or detection of the protein by antibodies. |
| Transcript-level evidence | Experimental evidence at transcript level | The existence of a protein has not been strictly proven but that expression data (such as existence of cDNA(s), RT-PCR or Northern blots) indicate the existence of a transcript. |
| Other | Protein inferred by homology | The existence of a protein is probable because clear orthologs exist in closely related species. |
|  | Protein predicted | Entries without evidence at protein, transcript, or homology levels. |
|  | Protein uncertain | The existence of the protein is unsure. |

### Integrating prediction outputs

The outputs from the independent prediction servers were parsed and integrated using Python 3.7. The source code is made available as a Jupyter Notebook to enable implementation on batch predictions from TMHMM, Phobius, PrediSi and SignalP^30,36,39,45^ for other species with minimal alteration (https://github.com/GundryLab/cirfess). A schematic of the inputs and the generated data structure is shown in Figure 1.

**Figure 1:**
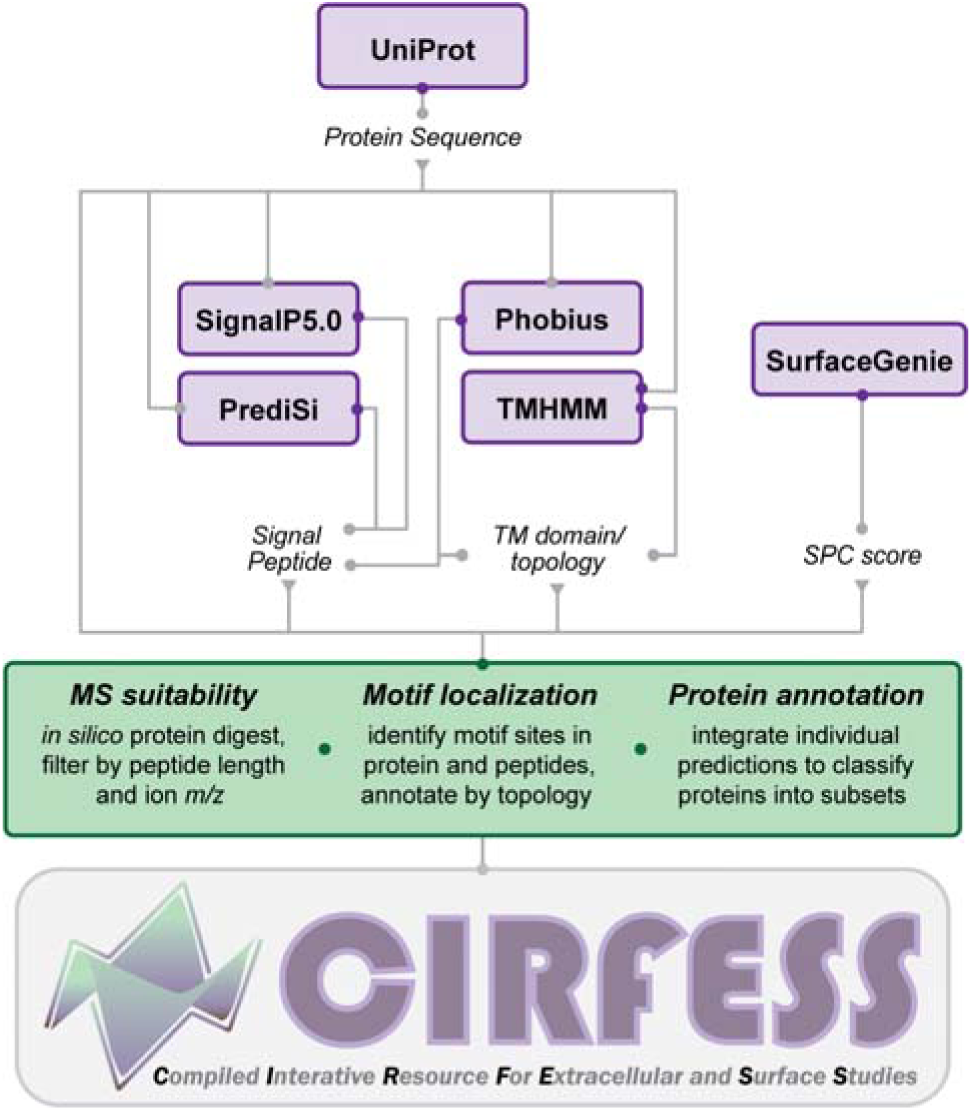
Schematic of the resources used (in purple) and the analyses performed (in green) for the construction of CIRFESS.

### Evaluating peptides, motifs, and topology

Protein sequences were digested *in silico* to generate a list of potential peptides using the canonical tryptic cleavage site, X[R/K] where X is not P, allowing for up to two missed cleavages. This list of peptides is subsequently annotated with the following information: (1) presence of motifs for relevant proteomic capture strategies; N[!P][S/T/C/V] for *N*-glycan based capture, C for cysteine-based capture, K for lysine-based capture, (2) topological information – which residues and motifs are predicted to be intracellular and extracellular, and (3) suitability for a standard bottom-up proteomic experiment – length > 5, *m/z* of 2+ or 3+ charge state peptide < 2000.

### CIRFESS Web application

A web application (CIRFESS) for accessing the data structure containing the parsed prediction outputs was developed in R^46^ using the Shiny package and is available at www.cellsurfer.net/cirfess. Source code is available at https://github.com/GundryLab/cirfess.

### Statistical Analysis

Chi-square analyses were performed using the *chisq.test* function in R (version 3.6.2). Students t-test were performed using the *t.test* function in Excel.

## Results and Discussion

### Proteome-wide comparison of prediction strategies

There are multiple sources of information to consider for evidence of surface or extracellular localization. Here we utilized transmembrane (TM) predictions^30,36^, signal peptide (SP) predictions^31,39^, and the Surface Prediction Consensus (SPC)^47^ score. To highlight the complementarity of these measures and justification for inclusion in this resource, we performed set analysis of the human proteome using the UpSetR^48^ web application. As shown in Figure 2A, the combination of these measures reveals distinct subsets of proteins. For instance, proteins which have a SP, but no TM domain (subsets b,f) are often considered to be the set of secreted proteins. For the set of proteins with predicted TM domains and SP, the SPC score is helpful for distinguishing between organelle membrane proteins (subset d) and cell surface proteins (subset a). Finally, proteins with an SPC score but not TM or SP may contain GPI-anchored proteins or those without a canonical SP (subset g).

**Figure 2:**
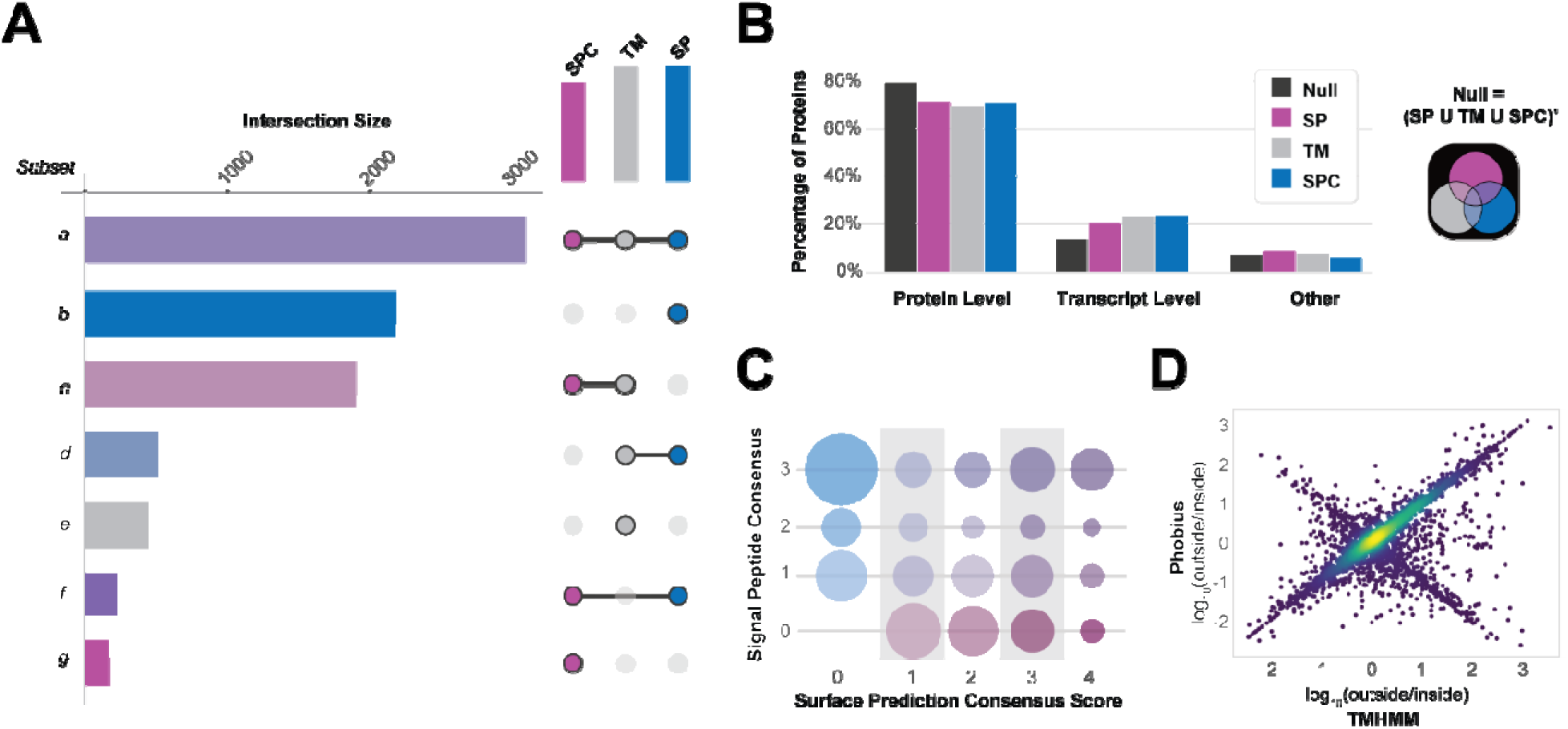
Contrasting views of the human proteome based on prediction strategies relevant for the cell surface proteome. (A) UpSet plot illustrating overlap in human proteins that are classified as containing a SP, TM domain, or SPC > 0. (B) Bar graph depicting the percentage of the human proteome with different levels of evidence, gathered from the “Protein existence” level listed for each accession number in UniProt, that are classified as containing a SP, TM domain, or SPC >0, or none of these features (Null). (C) Relationship between different levels of consensus for SP prediction and SPC score. Here, SP prediction consensus was calculated in a manner analogous to SPC score, where the number of positive SP predictions from SignalP, PrediSI, or Phobius was summed to generate a consensus score ranging from 0 to 3. (D) Plot depicting the log_10_ ratio of extracellular to intracellular residues predicted by TMHMM and Phobius highlighting that the opposite orientation is predicted for a subset of proteins.

To assess the level of experimental data that currently exists for cell surface proteins, we considered the “Protein existence” annotation within UniProt. For these and further analyses, we compared proteins with positive SP predictions, positive TM predictions, or those with SPC scores > 0, with proteins that were negative for all three analysis – which we term the ‘Null set’ of proteins (shown visually in Figure 2B). Compared to the Null set, all three classes of proteins have a lower percentage of members with protein-level evidence, 79% for Null proteins compared to 71%, 69%, and 71% for SP, TM, and SPC proteins, respectively (Figure 2B). The difference between the observed frequencies of the different levels of evidence were significant between Null and each other subset of proteins (SP, TM, and SPC), as revealed by Chi-square testing (with p-values of <2.2 × 10^−16^ for each test). Though mass spectrometry is not the only source of protein-level evidence for UniProt (see Methods), a potential explanation for this discrepancy is a statistical difference in the number of MS-suitable peptides between these subsets, as revealed by Student’s t-test summarized in Table 2. Nevertheless, this analysis highlights the need for further experimental investigation of the cell surface proteome as this class is less well-represented by experimental evidence than other subproteomes.

**Table 2:** Summary of the average number of peptides per protein that are “ok for MS” in different subsets of proteins. The t-test p-values were calculated by comparing the distribution to the Null-set of numbers of peptides (with the corresponding amount of numbers of missed cleavages).

|  | Null |  |  | SP |  |  | TM |  |  | SPC |  |  |
| --- | --- | --- | --- | --- | --- | --- | --- | --- | --- | --- | --- | --- |
| Missed Cleavages | 0 | ≤ 1 | ≤ 2 | 0 | ≤ 1 | ≤ 2 | 0 | ≤ 1 | ≤ 2 | 0 | ≤ 1 | ≤ 2 |
| Mean # of Peptides / protein | 33.1 | 87.5 | 146.4 | 28.3 | 70.5 | 112.3 | 26.2 | 65.0 | 103.4 | 28.1 | 69.8 | 111.0 |
| Median # of Peptides / protein | 25 | 66 | 109 | 20 | 50 | 79 | 19 | 48 | 76 | 20 | 50 | 79 |
| t-test p-value (compared to Null-set) | - | - | - | $6 \times 10^{-23}$ | $1 \times 10^{-41}$ | $2 \times 10^{-58}$ | $2 \times 10^{-47}$ | $8 \times 10^{-75}$ | $1 \times 10^{-97}$ | $2 \times 10^{-23}$ | $4 \times 10^{-42}$ | $4 \times 10^{-59}$ |

Another benefit of integrating these disparate predictions into a single analysis is revealed by looking at examples for which they do and do not agree. For example, stratifying proteins with positive SP predictions (6026 proteins) by the number of algorithms for which it was positive reveals that slightly over half (3192, 53%) are predicted by all 3 algorithms. Here, SP prediction consensus was calculated in manner analogous to SPC score, where the number of positive SP predictions from SignalP, PrediSI, or Phobius was summed to generate a consensus score ranging from 0 to 3. Plotting the number of positive SP predictions against SPC score reveals a positive relationship between SPC score and number of positive SP predictions for proteins with SPC score >0 (Figure 2C). However, it also reveals that the majority of proteins with three positive SP predictions has an SPC score of 0 (1648 of 3192, 51.6%). This suggests that secreted proteins may contain signal peptide sequences that are easier to recognize by prediction algorithms than proteins translocated through the membrane.

Focusing on TM proteins, TMHMM and Phobius predict 5353 and 5471 proteins with TM domains, respectively, with 4846 proteins in common and 1132 proteins unique to a single prediction strategy (507 and 625 in TMHMM and Phobius, respectively). While overall there is strong consensus between the two algorithms for predicting which proteins contain TM domains, the number of TM domains predicted differs for 1306 out of the 4846 commonly predicted proteins. Furthermore, the opposite membrane orientation was predicted for a subset of proteins, visualized by plotting the log_10_ ratio of the predicted extracellular to intracellular residues (Figure 2D). Altogether, these analyses demonstrate the value of integrating data from multiple sources and reveal that no single feature is sufficient to comprehensively predict the set of cell surface and extracellular proteins.

### Motif coverage of extracellular and surface predicted proteins

As prediction strategies alone are insufficient to define the set of proteins localized to the cell surface and extracellular space, experimentation is required. To aid in the selection of proteomic strategies that are likely to produce the desired coverage of the cell surface proteome, the SP, TM, and SPC analyses described above were integrated with *in silico* analyses designed to predict which proteins would generate tryptic peptides likely to be detectable by electrospray MS, and of those, which are expected to be captured by application of commonly used biorthogonal enrichment strategies targeting *N*-glycans and lysines^23–25,27,49–54^. We also considered cysteines as they are enriched in surface proteins compared to nonsurface proteins^40^ and numerous affinity reagents are available for targeting cysteines^55^, although this is not yet a widely described approach for cell surface proteins. Important for the *N-*glycan approach, although strategies that specifically enrich peptides from the extracellular space (glycan biotinylation is performed on cells with intact plasma membranes) provide an additional level of experimental evidence for surface localization, it is possible to capture *N-*glycopeptides from whole cell lysate. In this case, the MS-based evidence for a glycan modifying an asparagine within the consensus motif for *N*-glycosylation is proposed to serve as standalone evidence for surface localization. Canonically, the consensus motif has been described as NXS/T where X is any amino acid except proline. However, more recently, evidence for *N*-glycosylation has been put forth at NXC^56,57^ and NXV^58^. Here, we investigated the frequency of the various consensus motifs occurring in SP, TM, SPC and Null (meaning the protein contains no SP, TM, or SPC) sets of proteins. First, the probability of each motif occurring within the subset of proteins was calculated with respect to the amino acid frequencies. The expected frequencies based on amino acid compositions was consistent among the sets of proteins for each motif (0.26 ± 0.003 %, 0.18 ± 0.009 %, 0.09 ± 0.009 %, and 0.21 ± 0.019 % for NXS, NXT, NXC, and NXV respectively). Next, the observed frequency of each motif was calculated for each subset of proteins. The natural log of the odds ratio of observed to expected for each subset of proteins was calculated and plotted for each motif as well as the canonical (NXS/T) and complete consensus motifs (NXS/T/C/V) (Figure 3A). The results reveal that the NXS and NXT occur more frequently than expected and NXC and NXV occur less frequently than expected for SP, TM, and SPC proteins. Whereas NXS and NXT occur at about the expected rate for the Null set of proteins, NXC and NXV occur slightly above the expected frequency. While the complete consensus motif occurs more frequently than expected for SP, TM, and SPC proteins, the canonical motif demonstrates a much higher odds ratio, especially relative to the odds-ratio for the Null set. This analysis suggests that while the mere presence of the consensus motif provides some evidentiary weight to the localization of a surface protein – (1) it should not be considered conclusive, and (2) the canonical motif provides more meaningful information than the complete consensus motif. Based on these analyses, we elected to only consider the canonical motif as potential targets for *N*-glycan capture for subsequent analyses.

**Figure 3.**
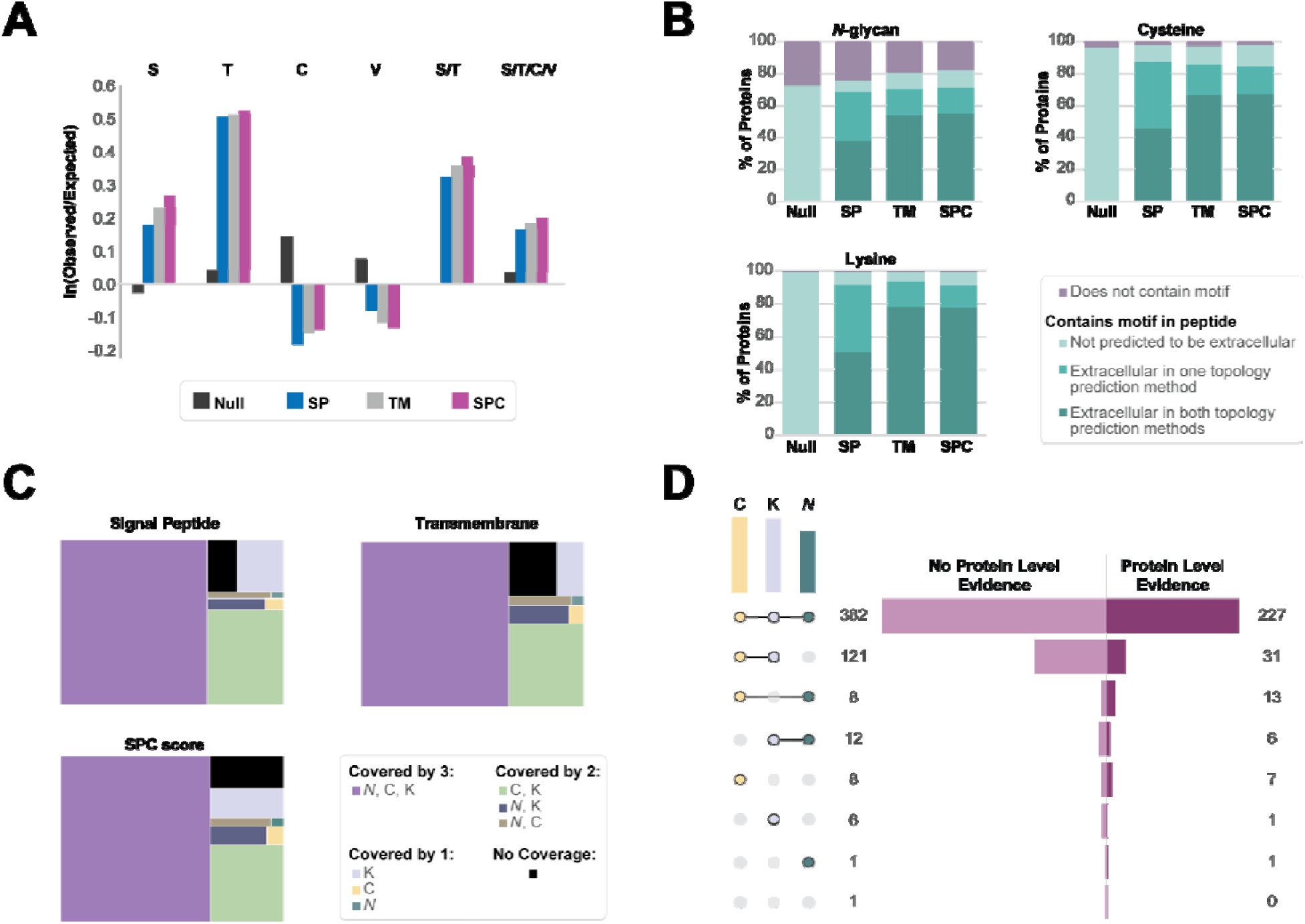
Results of CIRFESS analysis of the human proteome to assess predicted coverage provided by three common cell surface proteomic enrichment strategies. (A) The natural log of the odds ratio for observed-to-expected frequency of each permutation of the *N-*glycan consensus motif along with the canonical (S/T) and complete (S/T/C/V) consensus motif. (B) The expected coverage of the different subsets of proteins for each enrichment strategy broken down by which proteins have peptides with predicted extracellular motifs by one or both TM prediction methods. (C) The makeup of SP, TM, and SPC score proteins based on the overlapping coverage of the three individual enrichment strategies. (D) The set of human GPCRs based on expected coverage for enrichment strategy and level of evidence in UniProt.

By integrating the topology information provided by the TM predictions with the locations of motifs within proteins, we estimated the coverage that each capture strategy would provide for each subset of the proteome (Null, SP, TM and SPC). For this analysis, proteins were categorized based on whether they contained the relevant motif and whether the motif was in a region determined to be extracellular by one or both TM prediction strategies. The percentage of proteins for which a predicted extracellular motif was located within an MS-suitable peptide was recorded (Figure 3B). This analysis revealed that while 72% of Null proteins contain a consensus motif for *N-*glycosylation, none of the glycopeptides are predicted to be in the extracellular domain. In contrast, of the SPC proteins which contain the consensus motif, 86% of those proteins contain at least one peptide contains the consensus motif within the predicted extracellular domain. These results were further summarized by calculating the percentage of each subproteome that is predicted to be covered by each or multiple capture strategies (Figure 3C). Overall, querying the results from this analysis provides a strategy for investigators to rapidly interrogate the human proteome to determine which experimental strategy is most likely to be useful to address their biological question. In summary, 66.4 ± 0.4% of SP, TM and SPC proteins are likely to be captured by any of the three strategies, 17.3 ± 1.8% are detectable by cysteine or lysine capture, but not detectable by *N-*glycan strategies, 7.4 ± 1.4% are detectable by a single strategy, and 5.7 ± 1.2% are not detectable by any of the three strategies considered here. The identity of the proteins within each classification are provided in Supplemental Table 1A-C and these results provide actionable data related to high interest targets. For example, of the 825 human G-protein coupled receptors (GPCR), a striking 65.3% lack protein-level evidence within UniProt. Of these, all but one are predicted to be captured by at least one enrichment strategy and 70.9% of them are predicted to be captured by all three strategies. Supplementary Table 1D contains the identity of the GPCR proteins and which enrichment strategies are predicted to capture them.

### Critical Considerations

Results from the current implementation of CIRFESS are limited to human proteins digested with trypsin and the resulting peptides are detectable in the 2+ or 3+ charge state. These criteria were selected based on common implementation of bottom-up proteomic methods. However, all source files and code are publicly available in the Github repository and a user-specific version of CIRFESS could be generated, requiring minimum alteration to change the *in silico* digestion strategies or criteria for MS-compatible peptide filtering. Implementation on other species would require submission of proteins to the individual prediction servers, but the source code includes scripts to parse and integrate the generated output files. Another critical assumption is related to the *N-*gycan capture strategy where detection depends on the glycosite being occupied by a glycan which is sensitive to the oxidation strategy applied (*e.g.* cis diols for meta-periodate^59^). Currently, as it is not possible to predict which sites will be occupied with specific glycan structures, the peptides predicted to be observable by this strategy should be considered with this caveat in mind. It is possible that post-translational modifications may interfere with the digestion, capture, ionization, and identification of peptides in any of the strategies, and therefore experimental observations may not be fully predictable by this bioinformatics approach. Among the post-translation modification which may interfere with cysteine-based capture are disulfide bridges, which were ignored in this analysis, but a reduction step could be included prior to labeling in such an approach. Moreover, for enrichment strategies which use cleavable linkers, residual portions of the linker that remain after cleavage will increase the mass of the resulting peptide. However, the 2000 *m/z* range used here for predicting detectable peptides should accommodate most commonly used reagents. Finally, it may be beneficial to combine the results from CIRFESS analysis with predictions for peptide detectability^60–62^ or proteotypicity^63^ to better inform the set of peptides which are most likely to be observable or informative.

## Conclusion

CIRFESS is a web-based tool designed to accelerate cell surface proteome studies by eliminating the need to query each bioinformatics source separately and integrating disparate features into a single streamlined resource and output. Within the CIRFESSS interface, users are able to perform single and batch querying of protein accession numbers to extract protein-level and peptide-level annotations as well as information about numbers of motifs and motif-containing peptides. Results may be queried for proteins or protein classes of interest to inform the design of the next experiment. We anticipate that CIRFESS will be broadly applicable for multiple applications across a broad range of biology and disease studies. While there still exist significant technical challenges associated with the implementation of these technologies, particularly on sample-limited systems, these analyses suggest that acquiring protein-level evidence for the majority of predicted cell surface proteins is a matter of applying the right technology to a relevant biological system. Overall, we expect CIRFESS will promote the rational selection of the most apt cell surface proteomic methods and will inspire continued method development (*e.g.* cysteine-targeting) to enable detection of the human proteome not predicted to be accessible by established surface protein enrichment methods.

## Author Contributions

R.L.G. and M.W. conceived the study; R.L.G. supervised the study; M.W. and J. L. developed the algorithms and developed the web application; M.W. and R.L.G. analyzed data; M.W. generated figures; M.W. and R.L.G. co-wrote the manuscript; All authors approved the final manuscript.

## Acknowledgements

This work was supported by the National Institutes of Health [R01-HL126785, R01-HL134010, P20GM104320 (as a pilot grant award) to R.L.G.; F31-HL140914 to M.W.]; and JDRF [2-SRA-2019-829-S-B to R.L.G.]; Special thanks to Dr. Christopher Ashwood and Linda Berg Luecke for critical review of the manuscript and insightful discussions and Dr. David Tabb for helpful consultation. Funding sources were not involved in study design, data collection, interpretation, analysis or publication.

## Supplementary Information

**Supplemental Table 1.** A. Coverage of SP proteins. B. Coverage of TM proteins. C. Coverage of SPC Proteins. D. Coverage and evidence of G-protein coupled receptors.

